# The rise and the fall of a *Pseudomonas aeruginosa* epidemic lineage in a hospital

**DOI:** 10.1101/2020.09.21.307538

**Authors:** Marie Petitjean, Paulo Juarez, Alexandre Meunier, Etienne Daguindau, Hélène Puja, Xavier Bertrand, Benoit Valot, Didier Hocquet

**Author notes:** Didier Hocquet and Benoit are co-last authors and contributed equally to this work.

## Abstract

The biological features that allow a pathogen to survive in the hospital environment are mostly unknown. The extinction of bacterial epidemics in hospitals is mostly attributed to changes in medical practice, including infection control, but the role of bacterial adaptation has never been documented. We analyzed a collection of *Pseudomonas aeruginosa* isolates belonging to the Besançon Epidemic Strain (BES), responsible for a 12-year nosocomial outbreak, using a genotype-to-phenotype approach. Bayesian analysis estimated the emergence of the clone in the hospital five years before its opening, during the creation of its water distribution network made of copper. BES survived better than the reference strains PAO1 and PA14 in a copper solution due to a genomic island containing 13 metal-resistance genes and was specifically able to proliferate in the ubiquitous amoeba *Vermamoeba vermiformis*. Mutations affecting amino-acid metabolism, antibiotic resistance, lipopolysaccharide biosynthesis, and regulation were enriched during the spread of BES. Seven distinct regulatory mutations attenuated the overexpression of the genes encoding the efflux pump MexAB-OprM over time. The fitness of BES decreased over time in correlation with its genome size. Overall, the resistance to inhibitors and predators presumably aided the proliferation and propagation of BES in the plumbing system of the hospital. The pathogen further spread among patients via multiple routes of contamination. The decreased prevalence of patients infected by BES mirrored the parallel and convergent genomic evolution and reduction that affected bacterial fitness. Along with infection control measures, this may have participated in the extinction of BES in the hospital setting.

**Importance:** Bacterial pathogens are responsible for nosocomial outbreaks, but the sources of contamination of the hospitals are mostly unclear and the role of bacterial evolution in the extinction of outbreaks has never been considered. Here, we found that an epidemic strain of the pathogen *Pseudomonas aeruginosa* contaminated the drinking water network of a hospital due to its tolerance to copper and predatory amoeba, both present in the water pipes. The extinction of the outbreak occurred concomitantly with parallel and convergent genome evolution and a reduction in the size of the bacterial genome that correlated with the fitness of the pathogen. Our data suggest that pathogen evolution participated in the extinction of an outbreak in a hospital setting.

## Introduction

The epidemic curves of bacterial pathogens in hospitals are mostly wave-shaped, with a rapid increase in cases followed by a slower decline (1). During pathogen outbreaks in hospital settings, patients are infected directly or indirectly, either from the environment or from other patients. However, the exact source of the outbreak (*e.g.* a patient, a healthcare worker, the environment) is almost never documented and often limited to the identification of an index case due to the lag between contamination of the healthcare setting and detection of the pathogen in the index case (2, 3). Aside from antibiotic resistance, the biological features that allow a pathogen to enter and survive in the aggressive hospital environment are mostly unknown. However, genomic analyses are able to estimate the origin of an outbreak by calculating the time of emergence of the most recent common ancestor and the genetic features that could have favored its expansion (4, 5).

Whole-genome sequencing analysis has already been used to identify genetic elements gained, lost, or mutated during outbreaks of bacterial pathogens, but the phenotypic consequences of such mutations are rarely determined (6–8). Other studies have illustrated the difficulty of phenotypically confirming *in silico* predictions (9). Although the extinction of epidemics in hospitals is mostly attributed to infection control or more widely to changes in medical practice (10), the role of bacterial adaptation has never been considered.

We addressed this question by exploring the genotypic and phenotypic variations of an epidemic strain of *P. aeruginosa* that infected more than 250 patients over 12 years in a single hospital (2). *P. aeruginosa* is a ubiquitous Gram-negative species that exhibits such genomic plasticity that it allows its adaptation to a variety of niches, from the wild environment to that of disinfected hospitals and patients treated with antibiotics (11). This opportunistic pathogen is a leading cause of nosocomial infections, as well as chronic lung infections in cystic fibrosis (CF) patients (12). In addition to its high intrinsic resistance to a wide variety of antibiotics, *P. aeruginosa* can readily acquire resistance through mutations or horizontal transfer of resistance determinants. Thus, resistance to all currently available anti-pseudomonal antibiotics is frequent (13), justifying its inclusion in the list of serious threats by the CDC (14). *P. aeruginosa* is frequently responsible for outbreaks in the hospital setting, in which the source of contamination is mostly unknown or speculated, at best (3). Water points-of-use have often been incriminated in *P. aeruginosa* hospital outbreaks. However, sinks are themselves seeded by clones of human origin, blurring the tracks back to the source of hospital contamination (15).

We used WGS analysis and a genotype-to-phenotype approach of such a longterm outbreak of *P. aeruginosa* as a model to (*i*) date the true origin of the contamination of the hospital, (*ii*) identify the biological features that favored its installation, (*iii*) identify parallel and convergent evolution selected during the outbreak, and (*iv*) correlate the fitness of the strain with its genomic evolution.

## Results and discussion

### Population structure and most recent common ancestor of *P. aeruginosa* BES

Between May 1997 and April 2008, 276 patients were infected with the *P. aeruginosa* BES in the University Hospital of Besançon (Fig. 1A). We sequenced the full genomes of 54 isolates (retrieved from 54 patients) evenly distributed over time and considered to be representative of the outbreak (Table S1). *In silico* MLST showed the sequenced isolates to belong to the widespread clone ST395 (16). Phylogenetic analysis using 285 SNPs in the core genome showed that BES isolates fell into the two major clades, 1 and 2, with clade 1 being divided into four sub-clades, 1a to 1d (Fig. 1B). There was no clear link between the date of isolation or hospital ward and position in the phylogenetic tree (Table S1). Although no environmental sampling was made during the outbreak, clade 1b in the phylogenetic tree probably grouped isolates from patients contaminated from a common environmental reservoir. In addition, stair-shaped branches (*e.g.* clade 1c) indicated probable patient-to-patient transmission. Although the pulsed-field gel electrophoresis pattern of BES isolates was the same (2), WGS analysis showed two hidden parallel outbreaks simultaneously occurring in the same hospital (Fig. 1B), as already observed for other Gram-negative pathogens (17). The dating of emergence of the clone ST395 at the hospital required the calculation of the clock rate of our ST395 population, that was estimated to be 3.96 × 10^-6^ mutations per site per year by Bayesian analysis (95% highest posterior density, 3.08 × 10^-6^ - 4.96 × 10^-6^), which is higher than that estimated for the *P. aeruginosa* species (18). With the addition of an ST395 outgroup isolate retrieved in our hospital, but not belonging to BES, to the database (Table S1), Bayesian analysis dated the putative common ancestor that linked the ST395 isolates from our hospital to 1990 (95% confidence interval [CI] = 1987-1993), seven years before its first isolation from a patient in May 1997. Furthermore, using an external ST395 outgroup isolated from the hospital of Birmingham (United Kingdom), we estimated the emergence of clone ST395 at the hospital in 1978 (95% CI = 1970-1985) (Fig. 1B) during the creation of the water distribution network. The introduction of ST395 during the construction of a hospital was already suspected in the United Kingdom (3). This is consistent with the frequent isolation of the ST395 clone in water (19, 20) and the probable role of drinking water as a source of *P. aeruginosa* infection (21, 22). Overall, the ST395 clone has presumably contaminated the water network of the hospital before its opening but the pathogen further spread among patients by other routes of transmission (*e.g.* contact with the environment, other patients, or healthcare workers).

**Figure 1.**
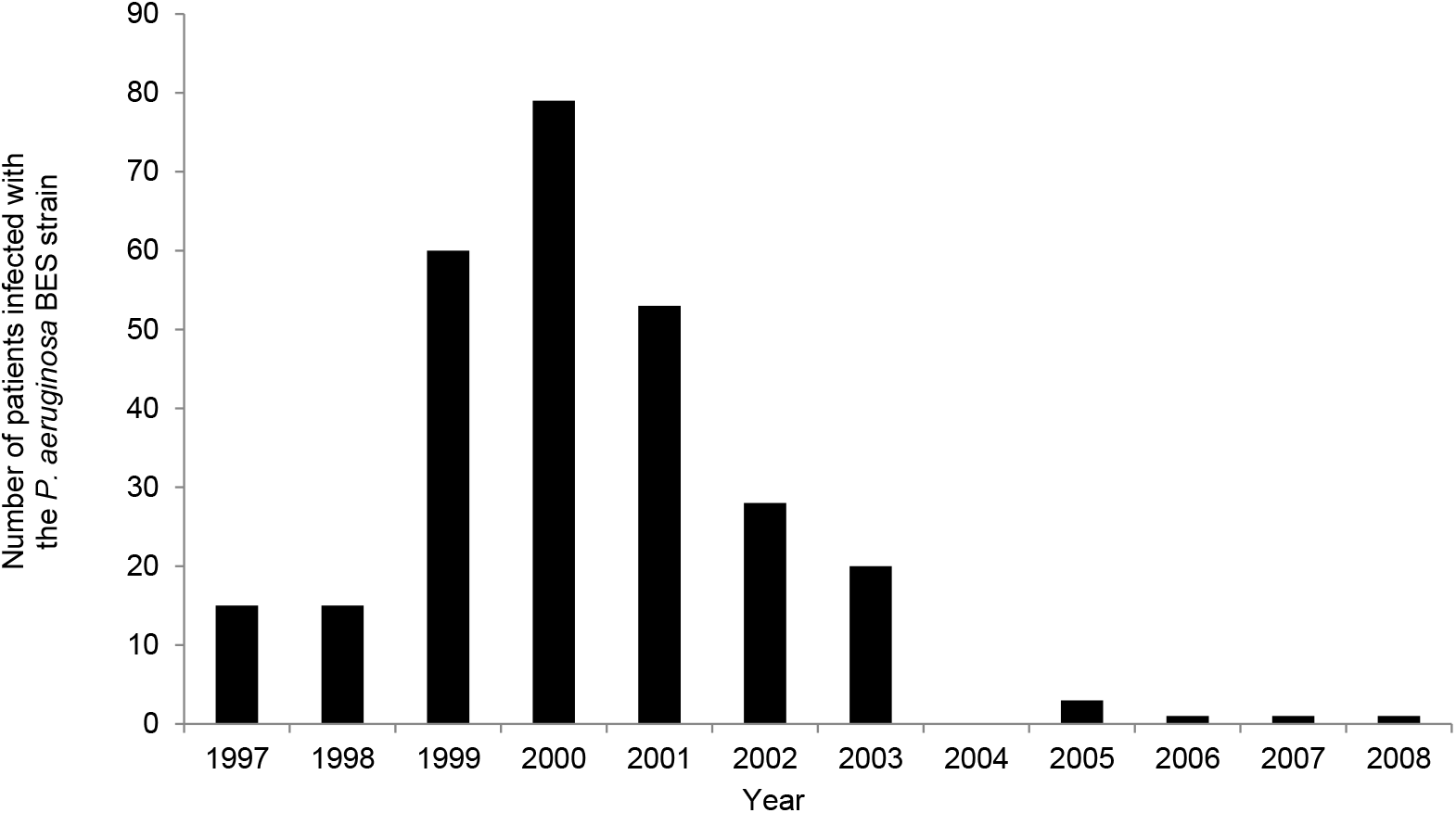
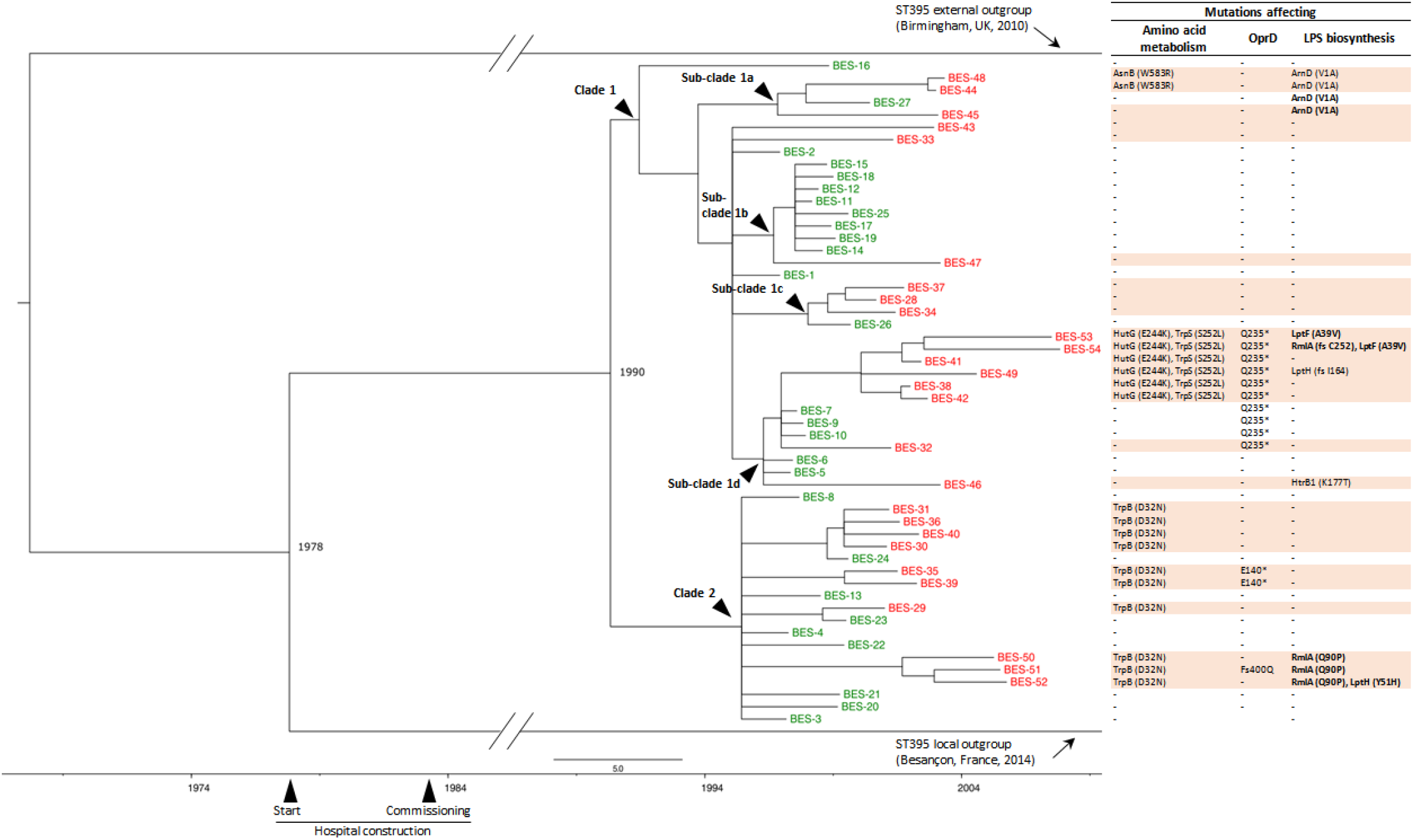
Temporal data and phylogenetic tree of the *P. aeruginosa* Besançon Epidemic Strain (BES) in a University Hospital. (*A*) Number of new patients contaminated by or infected with *P. aeruginosa* BES (belonging to the widespread clone ST395) by year during the 12 years of the outbreak in the University Hospital of Besançon (France). (*B*) The left panel shows the tree built with MrBayes using the following parameters: GTR substitution model and 1,000,000 cycles with the MCMC algorithm, with sampling of the chain every 1,000 cycles. We defined two major clades, 1 and 2, with clade 1 divided into four sub-clades, 1a to 1d. The isolates in green and red are early (before January 2001) and late (after January 2001) isolates, respectively. Isolates in black were outgroups. The right panel indicates non-synonymous mutations that affected amino-acid metabolism, OprD, or lipopolysaccharide (LPS) biosynthesis. Fs, frameshift. Mutations shown in bold indicate non-O:6 isolates.

### BES *P. aeruginosa* is tolerant to copper

The date of emergence of the common ancestor of ST395 (1978) led us to suspect the water distribution system of our hospital as the source of the outbreak. The water distribution and plumbing systems of our hospital are made of copper and are hostile environments for bacterial development, with nutrient-poor conditions, inhibitors (*e.g.* chlorine and copper-ions), and predators (free-living amoeba). We therefore postulated that BES could cope with those harsh conditions. Analysis of the BES genome sequences identified a 37-kb genome island (GI) that harbored an array of thirteen genes encoding copper transporters (Fig. S1) in all isolates of BES (23). This GI shared 99.7% identity with GI-7 of a *P. aeruginosa* ST308 copper-tolerant strain (24), also retrieved from water networks of hospitals. We phenotypically confirmed that the representative isolate BES-4 survived better than the *P. aeruginosa* reference strains PAO1 and PA14 (Fig. 2A) in a 150 μg/L copper (II) sulphate solution, correponding to the highest copper concentration retrieved in the drinking water of our hospital (24). We then tested the mutant strain BES-4ΔGI-7, deleted of the entire sequence of GI-7 by chromosomal recombination, under the same conditions to confirm the role of GI-7 in the tolerance of BES-4 to copper in solution. The deletion of GI-7 decreased the tolerance to copper of the BES strain relative to its wildtype parent, reducing it to that of the comparators PAO1 and PA14 (Fig. 2A). However, the tolerance to copper-ions under these conditions was not fully specific to GI-7, as the representative of the high-risk clone ST235 (isolate PA1646, without GI-7) showed copper tolerance similar to that of BES-4 (Fig. 2A). However, no ST235-specific determinants for tolerance to copper-ions have been identified (25). Interestingly, a sampling campaign in 2017 found that the same water distribution network was contaminated by a GI-7 containing ST308 clone (24). Overall, these results confirm that GI-7 can help waterborne *P. aeruginosa* (ST308, ST395) to tolerate copper in solution.

**Figure 2.**
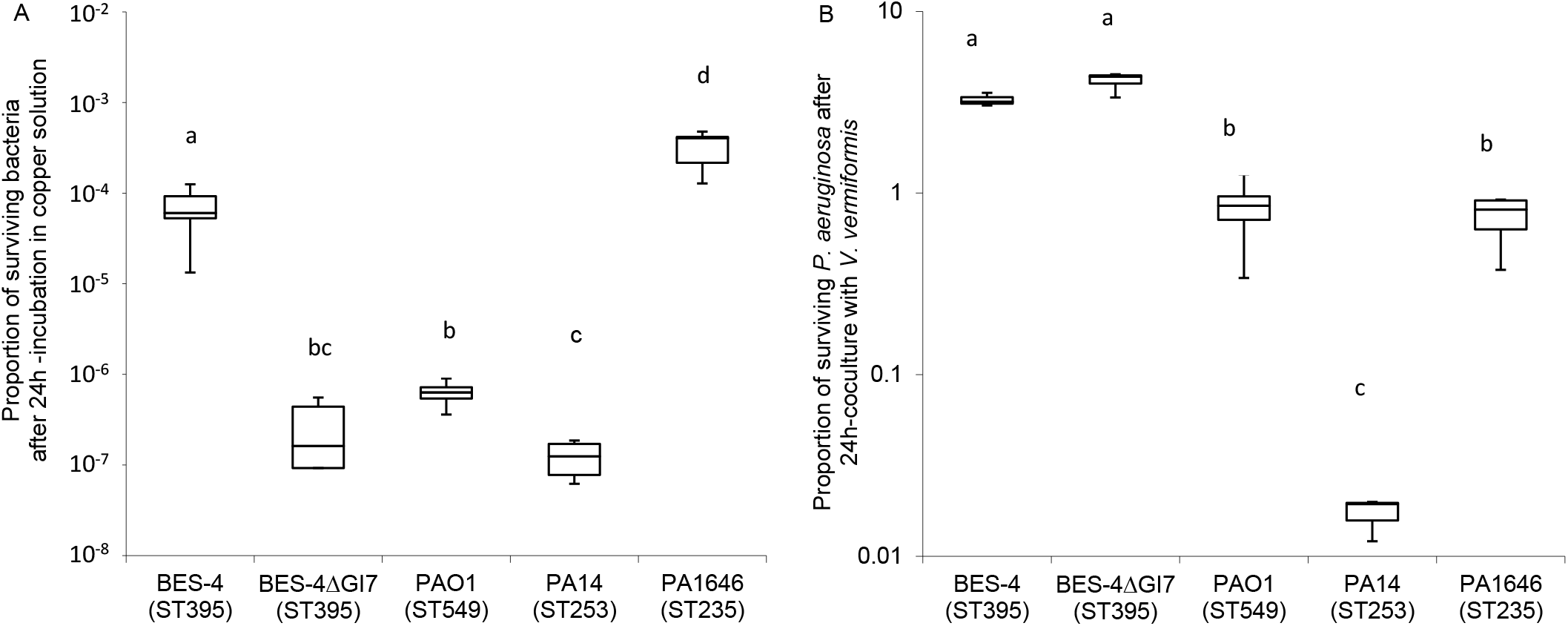
*P. aeruginosa* BES is tolerant to copper-ions and proliferates in the free-living amoebae *Vermamoeba vermiformis*. (*A*) Survival of *P. aeruginosa* BES-4, with or without GI-7, in a 150 μg/L copper (II) sulphate solution after 24 h of incubation shown on a decimal logarithm scale. Values are shown as the mean ratio ± SD (≥ 5 independent experiments). PCR experiments confirmed that the genomes of PAO1, PA14, and PA1646 (ST235) did not harbor GI-7. (*B*) The *P. aeruginosa* BES (ST395) proliferates in the free-living amoebae *Vermamoeba vermiformis.* The reference strain *V. vermiformis* 172A was inoculated with *P. aeruginosa* strains at a multiplicity of infection (MOI) of 5 and incubated for 24 h at 25°C. Values are shown as the mean ratio ± SD (3 technical replicates and ≥ 3 biological replicates). ANOVA (analysis of variance) with a Tukey HSD (honestly significant difference) test: a-b, a-c, a-d, b-c, b-d, and c-d are significantly different; *P* < 0.01. For the two panels, y-axis represent the ratio of the quantification on LB agar plates of the living cells at the en of the experiments and that of the initial inoculum.

### BES *P. aeruginosa* proliferates in free-living amoebae

Protozoans are predators for bacteria in water (26). *Vermamoeba (Hartmannella) vermiformis* is ubiquitous in drinking-water systems and resistant to commonly used disinfection methods (27). It has already been shown that a strain of *P. aeruginosa* isolated from the water of a hospital can resist *V. vermiformis* grazing (28). We thus tested the survival of BES-4 in the reference strain of *V. vermiformis* 172A versus that of the same controls used to test copper tolerance to further investigate whether this mechanism may contribute to the resistance of *P. aeruginosa* BES in the drinking-water network. BES better resisted the grazing of this protozoan than the comparators and was even able to proliferate inside the amoeba (Fig. 2B). Copper resistance has been implicated in the tolerance to amoeba, but the deletion of GI-7 had no effect on grazing resistance (29) (Fig. 2B). The resistance mechanism of BES to protozoan grazing is therefore yet unknown. Hence, protozoans were presumably responsible for the proliferation and propagation of BES in the water system and, as already demonstrated for *Legionella pneumophila*, the intracellular environment protected the bacterial pathogen from chlorine disinfection (30). However, such resistance to protozoan grazing could have broader consequences. Pathogens use related strategies for intracellular survival in both amoeba and human macrophages, which in turn share similar phagocytic mechanisms. Consequently, the coincidental evolution hypothesis suggests that virulence factors arose as a response to other selective pressures, such as predation, rather than for virulence *per se* (31).

Overall, the resistance to an inhibitor (copper) and a predator (free-living amoeba) presumably aided the proliferation and propagation of *P. aeruginosa* BES in the plumbing system during the building of the hospital, five years before its opening. This finding stresses the need for strict adherence to the procedures of commissioning water systems in healthcare premises (32).

### Biological functions altered during the 12-year spread of *P. aeruginosa* BES

After having identified the likely source of contamination of the hospital building and the associated biological features, we explored the evolution of BES during its spread. During their interaction with humans, bacterial pathogens can accumulate mutations, allowing them to better adapt to the host, evade the immune response, or increase their resistance to antibiotics (33). Our collection covered the 12-year epidemic and enabled the identification of parallel evolution of the same strain in multiple individuals. We then searched for biological features selected or lost during the outbreak. Variant calling retrieved 458 SNPs across the BES population, amongst which 220 were non-synonymous or present in the promoting regions of genes (Table S3). Among them, 68 mutations affected 55 genes encoding proteins for which the function was known. We compared the proportion of altered functions in early and late isolates and identified four major functions that were affected (χ^2^ test, *P* < 0.05): amino-acid metabolism, antibiotic resistance, lipopolysaccharide (LPS) biosynthesis, and regulation.

Specific modifications occurred in genes involved in amino-acid metabolism. This was particularly observed in late isolates. The genes *hutG* and *trpS* were both mutated in the last six isolates of clade 1d, whereas the late isolates of clade 1a harbored mutated *asnB,* and those of clade 2 mutations in *trpB* (Fig. 1B). It has been predicted that mutations in *hutG* and *asnB* impair the metabolism of histidine and asparagine, respectively, and that *trpS* and *trpB* mutants contribute to the deficient synthesis of tryptophan (34). Surprisingly, substitution D32N in TrpB independently occurred in late isolates of clade 2 (Fig. 1B). Convergent evolution towards auxotrophy has been frequently described in CF isolates as a result of niche specialization (35), presumably leading to an epidemiological dead end (36). However, the mutations in amino-acid metabolism did not change the growth of the isolates of BES in minimal media containing asparagine or histidine as the sole carbon source (Fig. S2). This may be due to compensatory mutations in unannotated genes or the existence of alternative pathways for amino-acid metabolism.

The biosynthetic pathway of LPS is encoded by several genes responsible for the biosynthesis of lipid A, O-antigen, and transport towards the outer membrane (37). Eleven isolates, of which 10 were late isolates, displayed parallel evolution affecting genes known to be involved in LPS biosynthesis, either in O-antigen assembly (*rmlA*), LPS export (*IptF, IptH*), or lipid A biosynthesis (*htrB1, arnD*) (Fig. 1B). Serotyping confirmed that most (7 of 11) of these isolates were polyagglutinable, resulting in an altered LPS. This contrasted with all the other isolates of BES, for which serotype O:6 was found (Fig. 1B). LPS structures can be modified in response to environmental pressures (*e.g.* divalent cations, antimicrobial peptides) and surface interactions. Modification of the LPS of *P. aeruginosa* is an important factor in the adaptation of this pathogen to chronic infection. Hence, reduced LPS immunostimulatory potential, due to lipid A modification (in isolates BES-27, −44, −45, −46, and −48; Fig. 1B), may contribute to evasion of the immune system and thus the survival of the pathogen over the course of a chronic infection (38). However, LPS extracted from all the isolates of BES stimulated human peripheral blood mononuclear cells with the same intensity (Fig. S3).

### Evolution of antibiotic resistance during the spread of *P. aeruginosa* BES

Early isolates of BES were classified as multidrug-resistant (39), with decreased susceptibility to aminoglycosides (gentamicin, tobramycin), fluoroquinolones (ciprofloxacin), and β-lactam compounds (40). Such decreased susceptibility may be explained by chromosomal mutations in resistance-associated genes. Indeed, early isolates harbored *ant(2”)-Ia*, coding for a 2”-aminoglycoside nucleotidyl-transferase that is involved in aminoglycoside resistance. Moreover, β-lactam resistance can be explained by observed mutations in the regulating genes *ampR* (M1L in AmpR) and *ampD* (A134V and T139M in AmpD), responsible for increased production of the cephalosporinase AmpC and alterations in *mexR* (H107P in MexR) a regulator of the multi-drug efflux pump MexAB-OprM. Finally, all isolates also displayed canonical mutations in the quinolone-resistance determining regions of GyrA (T83I) and ParC (S87L), explaining the increased resistance to fluoroquinolones. BES did not acquire foreign resistance determinants during the outbreak. Sequence analysis of three early and 10 late isolates showed three different mutations in *oprD* (Fig. 1B) that correlated with higher resistance to imipenem (Welch Two Sample Test, *P* = 2.5 x 10^-5^). Resistance to carbapenem via OprD mutation, certainly favored by antibiotic treatment, has been shown to enhance fitness *in vivo* and to facilitate mucosal colonization and dissemination to the spleen (41).

### Modulation of *mexAB-OprM* overexpression

The resistance-nodulation-division efflux pump MexAB-OprM is expressed at basal levels by wildtype cells and regulated by a complex regulatory network of at least 10 direct or indirect regulators (42) (Fig. 3A). Early isolates displayed an alteration of the DNA-binding domain of the repressor protein MexR (H107P, substituting the polar histidine with the non-polar proline). This resulted in the overexpression of *mexB* relative to that of the PAO1 reference strain (Fig. 3B) and contributed to enhanced resistance to the MexAB-OprM substrates aztreonam and ciprofloxacin (Fig. S4) (43). Other mutations in MexAB-OprM-regulating regions arose independently during the spread of BES and were found in 19 of the 27 late isolates. Hence, 12 isolates of BES carried non-synonymous mutations in *mexR*, *cpxR*, or *nalD*, and seven had mutations in the promoter region of the *mexAB-oprM* operon (Fig. 3A). We verified the impact of these mutations on the expression of the efflux pump by measuring *mexB* expression in all isolates of BES by RT-qPCR. Almost all the isolates (29 of 31, 94%) that had the same mutations in the *mexAB-oprM* regulation pathway as BES-1 overexpressed *mexB* (median value, 4.3; Fig. 3B). Ten late isolates with a H107P MexR background had undergone three types of mutations in the regulator CpxR (Fig. 3A), leading to an intermediate level of *mexB* expression (median value, 3.0; Fig. 3B). Moreover, 11 isolates harbored mutations in the *mexA*-*mexR* intergenic region (MexR binding site, and *mexA* RBS region) or had a P107S substitution in MexR (returning a polar amino acid-serine - to position 107), expressed *mexB* at a level similar to that of the reference strain PAO1 (median value, 0.7; Fig. 3B). Efflux expression, as measured by *mexB* expression, also decreased over time (Fig. 3C).

**Figure 3.**
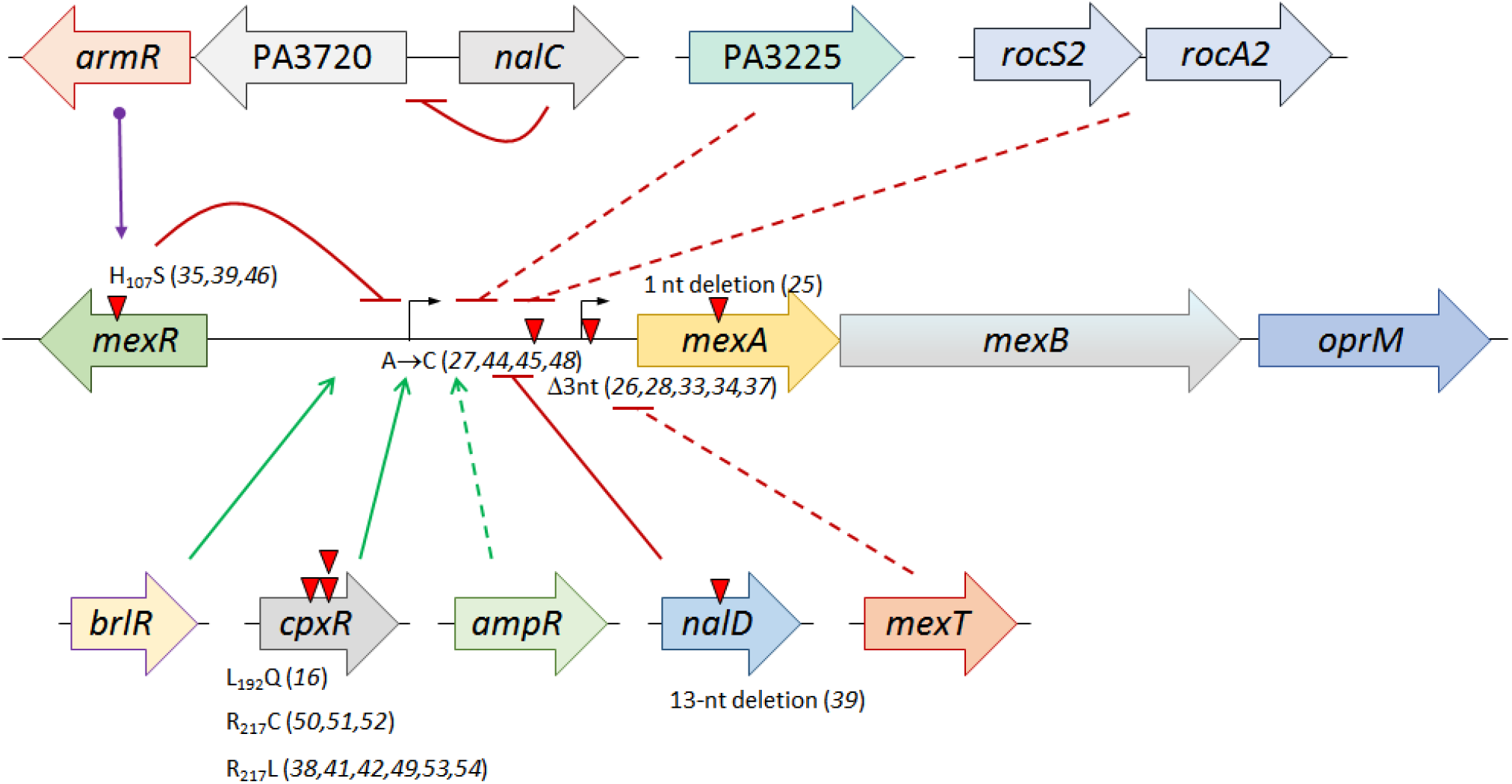
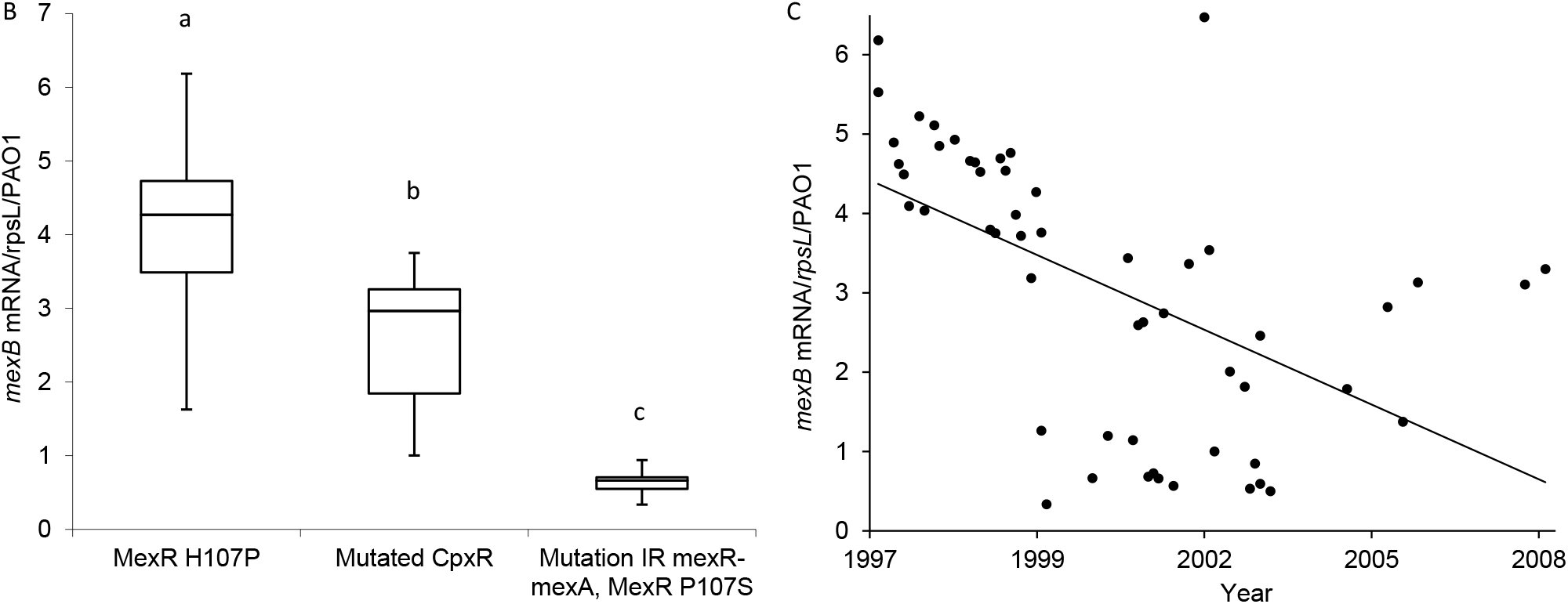
*P. aeruginosa* BES attenuates the overexpression of the RND efflux pump MexAB-OprM during its spread via parallel and convergent evolution. (*A*) Mutations affecting *mexAB-oprM* expression in *P. aeruginosa* BES in the regulatory network of *mexAB-oprM*. Local repressor MexR ensures basal expression of the *mexAB-oprM* operon. The repressor NalD directly binds to the proximal promoter of the operon. NalC indirectly modulates *mexAB-oprM* expression by repressing the expression of the *armZ* gene, the product of which acts as an anti-repressor of MexR. Mutations in these three repressors have been reported in *nalB*, *nalD*, and *nalC* mutants, respectively. Two direct activators of *mexAB-oprM* are CpxR and BrlR. Finally, the products of four other genes have been reported to have an indirect negative (*mexT, rocS1/2-A2, PA3225*) or positive (*ampR*) impact on *mexAB-oprM* expression (64). Red triangles indicate mutations occurring in regions involved in the regulation of *mexAB-OprM* expression, with the number of the affected BES isolates in parenthesis. (*B*). Expression of *mexB* in isolates of BES sorted by the type of mutation in the regulatory pathway of *mexAB-oprM.* ANOVA with a Tukey HSD test: a-b, a-c, and b-c are significantly different; *P* < 0.001. (*C*) Attenuation of *mexAB-oprM* overexpression in *P. aeruginosa* BES over time. Pearson correlation between *mexB* expression and the time of isolation: - 0.547 (*P* = 1.87 x 10^-5^).

Overall, seven distinct regulatory mutations attenuated the initial *mexAB-oprM* overexpression of BES (Fig. 3A). MexAB-OprM plays a role in antibiotic resistance, but can also export virulence determinants that allow *P. aeruginosa* to be invasive and cause infection (44). The attenuation of *mexAB-oprM* overexpression has been shown to reduce the invasiveness of the bacteria in a cell model (45). It is thus possible that convergent evolution decreasing the overproduction of this pump contributed to the persistence of BES in hospitalized patients. Moreover, the overexpression of *mexAB-oprM* reduces the survival of *P. aeruginosa* in the environment (46). As expected, attenuation of the overexpression of *mexAB-oprM* in late isolates reduced the level of resistance to aztreonam and ciprofloxacin (Fig. S4). The regulatory mutations in BES contrasted with the alterations in *mexAB-oprM* genes that are selected in *P. aeruginosa* populations in CF airways, leading to hypersensitivity to antibiotics (47). A functional MexAB-OprM pump is necessary for the full virulence of *P. aeruginosa* in cell models (48). However, its overproduction has no effect on the invasiveness of the pathogen, advocating against the enhanced survival of MexAB-OprM-overproducing mutants in protozoans (45). This fine-tuning of *mexAB-oprM* expression possibly allows a trade-off between antibiotic resistance, virulence, and survival in the environment. Such efflux modulation may then be important for survival and persistence of the clone in hospital setting.

### Insertion or deletion of gene blocks during the spread of BES

*P. aeruginosa* can evolve through the acquisition or loss of genome fragments (6). Accessory gene profiling showed clear differences in accessory gene content among isolates, with gene blocks specific to certain clusters (Table S4). For example, isolates BES-1 and BES-28, which were retrieved four years apart and belonged to different clades, independently underwent a ca. 130-kb deletion, including genes encoding quorum-sensing regulators (LasI, LasR, and RsaL) and the flagellar operon region *fli* (23). The late isolate BES-52 lost a 278-kb fragment that encompassed *hmgA*, of which deletion leads to an easily recognizable brown colony phenotype due to homogentisate accumulation. These melanogenic mutants are probably adapted to life in diverse bacterial communities in chronically-infected patients (49). Despite the relatively high plasticity observed throughout the outbreak, the decrease over time in the number of genes per genome was not statistically significant (Spearman correlation test, *P* = 0.064; Fig. S5). The epidemiology of *P. aeruginosa* BES at our hospital (Fig. 1A) supports the idea of emergence and extinction over time. As genome reduction can alter the bacterial growth rate (50), we evaluated the fitness of each BES isolate by coculturing them with an early representant of BES and measuring by qPCR the proportion of each competitor after 48 h of incubation. Coculture was preferred since it permits detection of fitness differences that are ≤ 1% per generation, undetectable through the assessment of growth rates in single culture (51). Indeed, the fitness of BES decreased over time (Fig. 4A) and correlated with the size of the genome (Fig. 4B). More specifically, non-O:6 isolates (*n* = 7) showed lower fitness in coculture than O:6 isolates (Welch two-sample test, *P* = 0.012).

**Figure 4.**
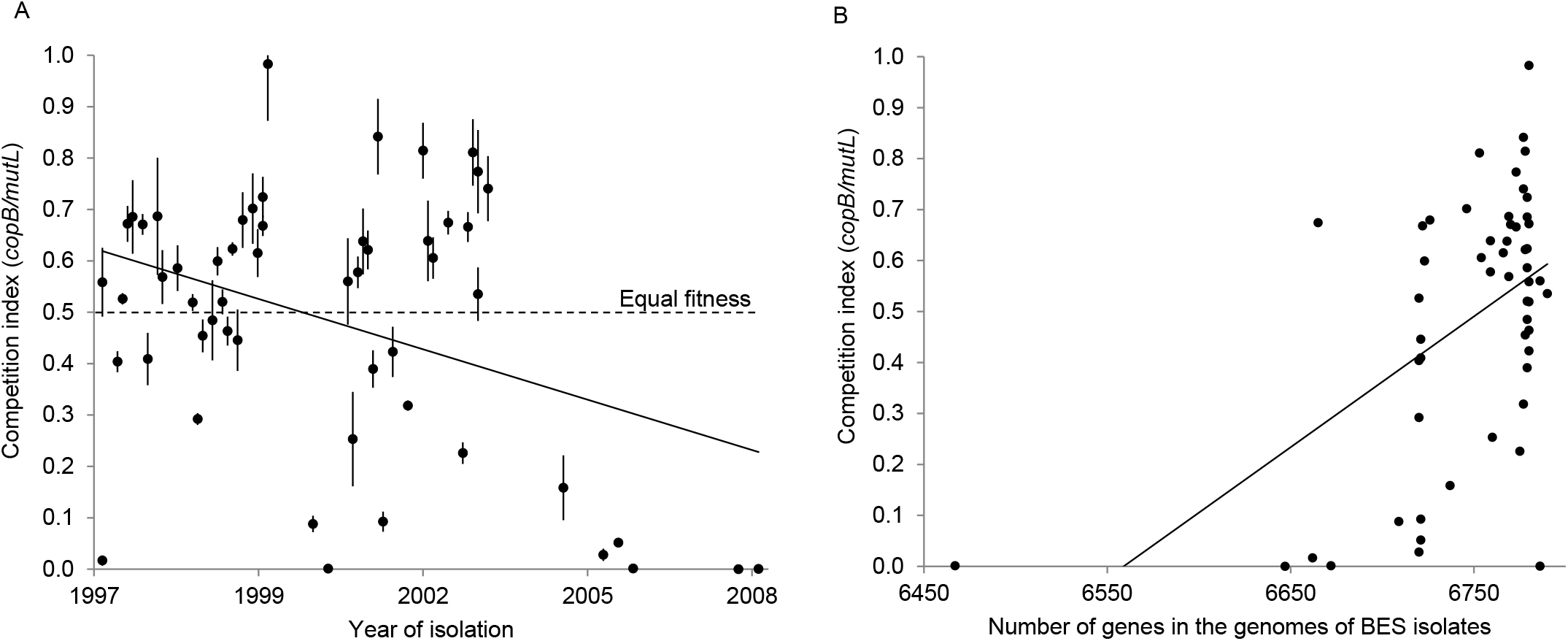
The fitness of *P. aeruginosa* BES decreases in relation to its genome size during a hospital-associated outbreak. (*A*) *P. aeruginosa* isolates of BES were cocultured in Luria-Bertani broth with the reference isolate *P. aeruginosa* BES-4ΔGI-7. After 48 h of incubation, we determined the *copB/mutL* ratio, which represents the proportion of the tested isolate of BES (harboring *copB* and *mutL*) in the co-culture relative to that of the isolate BES-4ΔGI-7 (harboring only *mutL*) and used it to calculate the competition index (CI), as described in Materials and Methods. Data are shown as the mean ± SD of two technical replicates, indicative of three biological replicates. The CI of each BES isolate is plotted against its time of isolation. The dotted line (CI = 0.5) represents an equal fitness. Pearson correlation test between the CI and the time of isolation: - 0.379 (*P* = 0.00465). (*B*) The fitness of *P. aeruginosa* BES correlates with the size of the genome. Pearson correlation test between the CI and the number of genes in the genome: 0.525 (*P* = 4.7 x 10^-5^).

### Limitations and strengths

This study had several limitations. Although we selected randomly distributed isolates throughout the period of the outbreak, the selection and inclusion criteria were not systematic and dependent on the availability of the isolates. Thus, the data presented here may be affected by a sampling bias that could have obscured epidemiological and evolutionary signals. This retrospective study did not allow water sampling at the time of emergence of the clone in the hospital (ca. 1978) that could have showed water contamination with BES and *V. vermiformis*. However, others have showed that *V. vermiformis* is endemic in hospital plumbing systems (52).

Here, we made use of a rare and valuable resource (*i.e.* longitudinally collected isolates from a variety of infection types in a single hospital from a period of more than a decade). While adaptation of human pathogens within a single niche has been described for evolution *in vitro*, very little research has focused on bacterial adaptive processes within the course of an outbreak. We attempted to validate *in silico* predictions by wet lab experiments. Hence, our genotype-to-phenotype approach correlated tolerance to copper, resistance to antibiotics, modulation of the expression of efflux pump, and bacterial fitness with genome content. However, sequence analysis predicted auxotrophy for tryptophan and impaired growth in histidine and asparagine, which were not confirmed *in vitro*. This highlights the need to control the functional effects of the mutations observed in WGS data.

### Conclusions

This whole-genome phylogenetic analysis of a *P. aeruginosa* outbreak suggests contamination of the water distribution system before the opening of the hospital with the widespread clone of *P. aeruginosa* ST395. Its resistance to copper and protozoans presumably helped the strain to survive in the water distribution system. Although BES has presumably contaminated the water network of the hospital before its opening, it further spread among patients by other routes of transmission. The prevalence of patients infected by the epidemic clone ST395 started to decrease in 2001, concomitantly with parallel and convergent evolution. Indeed, independent mutations affected the aminoacid metabolism pathways, led to the loss of porin OprD, and altered the LPS synthesis pathway. It also modulated the overexpression of the efflux pump MexAB-OprM, of which its fine-tuning may be important for survival and persistence of the clone. We found that the fitness of BES decreased over time, in correlation with genome length. The parallel adaptation of the ST395 isolates to patients may have decreased its resistance to the environment and thereby its patient-to-patient transmissibility. This, along with the implemented infection control measures, may have contributed to the extinction of this epidemic pathogen in a hospital setting.

## Materials and Methods

### Hospital setting and isolate collection

A prolonged outbreak of *P. aeruginosa* in the University Hospital of Besançon, a 1,300-bed teaching hospital in France, was investigated. The construction of its facilities began in 1978 and the main building was commissioned in 1983, with all pipes of the drinking-water installation being made of copper. All *P. aeruginosa* isolates retrieved from patients hospitalized between May 1997 and April 2008 were typed by pulsed field gel electrophoresis (PFGE) to assess the number of infected patients. Patients with *P. aeruginosa* isolates for which the pulsotype shared the same band profile with the index case (*n* = 276) were included in the cohort (2). None of them suffered from cystic fibrosis. Then, 54 isolates retrieved from 54 patients (BES-1 to BES-54) were selected for sequencing. They were evenly distributed over time and considered to be representative of the outbreak (Table S1). The first 27 isolates were recovered in the early stages of the outbreak (before January 2001) and the other 27 isolates were considered to be late isolates (Table S1).

### DNA sequencing and analysis

Bacterial DNA was isolated from overnight cultures on Mueller-Hinton agar using the Genomic-tip kit (Qiagen) and paired-end sequenced with Illumina NextSeq, at 2 x 150 bp. Reads were subsampled before assembly to limit the coverage to 80X. The contigs were built with SPAdes (53). Local and external outgroup isolates of *P. aeruginosa* of the same sequence type (ST) were used to root the phylogenetic tree (Table S1) (3). We previously sequenced BES-1, the first clinical isolate of BES identified in May 1997, using PacBio technology and used it as a reference (23). STs were determined *in silico* using the PubMLST database (https://pubmlst.org/) and the homemade pipeline pyMLST (https://github.com/bvalot/pyMLST). The BES pangenome was built by combining the BES-1 circular genome with non-redundant contigs from the other BES isolates using the Ragout tool (54). For this purpose, the assemblies of each BES isolate were aligned against the BES-1 genome using Mummer (55). Genes of the BES pangenome were detected using Prodigal and then annotated with BLAST (56, 57). Resistance and virulence genes were sought using ResFinder and the Virulence Factor DataBase, respectively (58, 59). The reads from all isolates were then aligned against the BES pangenome using BWA and the alignments used for further analysis (https://github.com/lh3/bwa). The presence or absence of each gene was tested using FeatureCounts (60). SNPs were identified using freebayes with a minimum coverage of 10 reads per isolate and a minimum quality of 30 (https://github.com/ekg/freebayes). Non-synonymous SNPs and SNPs in promoter regions (RBS and promoter sequences at −10 and −35) were further sorted. SNPs and gained or lost gene blocks were clustered in relation to the phylogenetic tree with MixOmics (61). IS insertion events were detected using panISa software (62). The boundaries of each potential IS were sought in the BES-1 genome to exclude chromosomal rearrangement before searching for them in the BES genomes to assess the length of the potential ISs. The presence of the potential IS was confirmed by PCR and sequencing with specific IS1 and IS2 primers (Table S2).

### Phylogeny and Most Recent Common Ancestor (MRCA) calculation

The 265 core genes of the BES isolates that harbored at least one SNP were concatenated for phylogenetic analysis, producing a 397,062-bp alignment with 285 SNPs. MrBayes software was used to build the maximum clade credibility tree (63). The isolation dates of the BES and outgroup isolates were used as calibration nodes to calculate the age of the MRCA.

### RT-qPCR experiments

*mexB* expression was assessed in all BES isolates by RT-qPCR as previously described (43) with specific primers for the housekeeping gene *rpsL*, and *mexB* (Table S2). After each assay, a dissociation curve confirmed the specificity of all PCR amplicons. The mRNA levels of *mexB* were normalized to that of the reference gene *rpsL* and expressed as a ratio to the levels in the reference strain *P. aeruginosa* PAO1. Based on previous results, all BES isolates with at least two-fold higher *mexB* expression than that in PAO1 were considered to be MexAB-OprM overproducers (43).

### Phenotype determination

Susceptibility to antipseudomonal compounds was tested using agar dilution, as recommended by the CLSI (64). We determined the serotype of all isolates with specific antiserum (Bio-Rad). Metabolic changes in BES isolates were assessed using procedures detailed in Fig. S2.

### Survival of the BES-4 isolate and its ΔGI-7 mutant in a copper solution

For all tested *P. aeruginosa* isolates, an overnight culture in Luria-Bertani broth (LBB, Bio-Rad) was diluted 1/200 into fresh LBB and further incubated at 37°C with gentle shaking until mid-log phase. The bacterial suspension was centrifuged and the pellet was washed once and resuspended in sterile deionized water. A 150 μg/L copper (II) sulphate solution (Sigma-Aldrich) was inoculated with approximately 10^8^ bacteria before incubation for 24 h at 22°C in the dark. The concentration of 150 μg/L was chosen as it was the highest value retrieved in the drinking water of our hospital (24). Initial and final bacterial concentrations were determined by plating appropriate dilutions on LB agar (Bio-Rad) and incubating overnight at 37°C. The ratios of the initial and final bacterial concentrations determined the survival rates. We repeated each experiment six times independently for all isolates. The reference strains of *P. aeruginosa* PAO1 (ST549), PA14 (ST235), and a clinical isolate of ST235 (strain PA1646)(24) were used as controls.

### Survival of *P. aeruginosa* in free-living amoebae

The survival and growth of the BES strain was evaluated in the free-living amoeba *Vermamoeba vermiformis* 172A, isolated from the drinking water installation of the Lausanne Hospital (Switzerland) (65). The protozoans were grown in PYNFH medium (LGC standards, Molsheim, France) as monolayers at 25°C before their harvest. *P. aeruginosa* strains were cultured on Mueller-Hinton agar (MHA) for 24 h at 37°C, harvested, and then suspended in PAS. Briefly, 1.5 x 10^5^ amoebae were incubated with *P. aeruginosa* at a MOI of 5 (five times more bacteria) for 4 h at 25°C. Non-internalized bacteria were then killed by a 2-h incubation with 0.3 mg.ml^-1^ amikacin. The co-cultures were washed three times with PAS to remove the antibiotic and further incubated in PYNFH for 24 h at 25°C. We then lysed the amoebae with 0.4% Triton X100 for 30 min at 25°C. Appropriate dilutions of the initial and final bacterial inoculum were plated on MHA plates to calculate the proportion of surviving bacterial cells.

### Chromosomal deletion of the genomic island GI-7

The GI-7 region was deleted from isolate BES-4 using overlapping PCR and homologous recombination detailed elsewhere (24).

### BES coculture

Mid-log phase BES isolates were co-cultured in LBB at a 1:1 ratio against the mutant BES-4ΔGI-7 as a reference. We extracted and purified the DNA (QIAamp DNA Mini Kit, Qiagen) from 48-h co-cultures. DNA concentrations were determined using QUBIT 2.0 and adjusted to 0.1 ng/μl. The *mutL* (present in all isolates) and *copB* (specifically absent from BES-4ΔGI-7) genes were quantified using specific primers and Taqman probes against standard curves (Table S2). qPCR was performed on a 7500 Fast Real Time PCR system and the results analyzed using 7500 software v 2.3 (Applied Biosystems). The competition index (CI, proportion of the tested BES isolate in the co-culture after 48 h of incubation) was calculated as the ratio of the concentrations of *copB* and *mutL.* In other words, a CI of 0.5 was neutral. The CI of BES-4 against BES-4ΔGI-7 was 0.53 ± 0.01, indicating that the deletion of GI-7 did not alter bacterial fitness.

## Data availability

All genomic data were deposited in NCBI under the project accession number PRJNA399056. Biosample details are given in the Table S1.

## Supplemental material

Figures S1 to S5

Tables S1 to S4

## Acknowledgments

We thank Marilou Bourgeon for her technical support and Catherine Llanes and Thilo Köhler for helpful discussions. We also thank Dr Beryl Oppenheim for the gift of a ST395 isolate from the Queen Elizabeth Hospital, Birmingham, UK. Computations were performed at the Mésocentre de Calculs de Franche-comté (Besançon, France).

X.B., B.V., and D.H. designed the research; M.P., P.J., A.M., H.P., and B.V. performed the research; B.V. contributed new reagents/analytical tools; M.P., P.J., E.D., B.V., and D.H. analyzed the data; and M.P., P.J., X.B., B.V., and D.H. wrote the paper.

This work was supported by a grant from the Région de Franche-Comté and from the University Hospital of Besançon, France (API 3A – Promes 2013) to M.P., the French “Ministère de l’Enseignement Supérieur et de la Recherche to P.J., and the Région Bourgogne Franche-Comté to H.P. The funders had no role in the study design, data collection or analysis, decision to publish, or preparation of the manuscript. Ethical approval was not required for this study.

